# Cryo-EM structures of α-synuclein filaments from Parkinson’s disease and dementia with Lewy bodies

**DOI:** 10.1101/2022.07.12.499706

**Authors:** Yang Yang, Yang Shi, Manuel Schweighauser, Xianjun Zhang, Abhay Kotecha, Alexey G. Murzin, Holly J. Garringer, Patrick W. Cullinane, Yuko Saito, Tatiana Foroud, Thomas T. Warner, Kazuko Hasegawa, Ruben Vidal, Shigeo Murayama, Tamas Revesz, Bernardino Ghetti, Masato Hasegawa, Tammaryn Lashley, Sjors H.W. Scheres, Michel Goedert

**Author notes:** These authors jointly supervised this work: Sjors H.W. Scheres, Michel Goedert. **Corresponding Authors** Correspondence to Sjors H.W. Scheres and Michel Goedert.

## Abstract

Parkinson’s disease (PD) is the most common movement disorder, with resting tremor, rigidity, bradykinesia and postural instability being major symptoms (1). Neuropathologically, it is characterised by the presence of abundant filamentous inclusions of α-synuclein in the form of Lewy bodies and Lewy neurites in some brain cells, including dopaminergic nerve cells of the substantia nigra (2). PD is increasingly recognised as a multisystem disorder, with cognitive decline being one of its most common non-motor symptoms. Many patients with PD develop dementia more than 10 years after diagnosis (3). PD dementia (PDD) is clinically and neuropathologically similar to dementia with Lewy bodies (DLB), which is diagnosed when cognitive impairment precedes parkinsonian motor signs or begins within one year from their onset (4). In PDD, cognitive impairment develops in the setting of well-established PD. Besides PD and DLB, multiple system atrophy (MSA) is the third major synucleinopathy (5). It is characterised by the presence of abundant filamentous α-synuclein inclusions in brain cells, especially oligodendrocytes (Papp-Lantos bodies). We previously reported the electron cryo-microscopy (cryo-EM) structures of two types of α-synuclein filaments extracted from the brains of individuals with MSA (6). Each filament type is made of two different protofilaments. Here we report that the cryo-EM structures of α-synuclein filaments from the brains of individuals with PD, PDD and DLB are made of a single protofilament (Lewy fold) that is markedly different from the protofilaments of MSA. These findings establish the existence of distinct molecular conformers of assembled α-synuclein in neurodegenerative disease.

---

A causal link between α-synuclein assembly and disease was established by the findings that missense mutations in *SNCA* (the gene that encodes α-synuclein) and multiplications (duplications and triplications) of this gene give rise to inherited forms of PD and PDD (7,8). Some mutations also cause DLB (8,9). Sequence variation in the regulatory region of *SNCA* is associated with increased expression of α-synuclein and a heightened risk of developing idiopathic PD, which accounts for over 90% of cases of disease (10). Both inherited and idiopathic cases of PD, PDD and DLB are characterised by the presence of abundant Lewy bodies and Lewy neurites in central and peripheral nervous systems (2).

α-Synuclein is a 140-amino acid protein, over half of which (residues 7-87) consists of seven imperfect repeats, which are lipid-binding domains (11). They partially overlap with a hydrophobic region (residues 61-95), also known as the non-β-amyloid component (NAC) (12), which is necessary for the assembly of recombinant α-synuclein into filaments (13). The carboxy-terminal region (residues 96-140) is negatively charged and its truncation results in increased filament formation (14). Upon assembly, recombinant α-synuclein undergoes conformational changes and takes on a cross-β structure that is characteristic of amyloid (15,16). The core of α-synuclein filaments assembled from recombinant protein *in vitro* extends from approximately residues 30-100 (17).

Seeded assembly of α-synuclein, propagation of inclusions and nerve cell death have been demonstrated in a variety of systems (18-21). Assemblies of α-synuclein with different morphologies display distinct seeding capacities (22,23). Indirect evidence has also suggested that different conformers of assembled α-synuclein may characterise disorders with Lewy pathology and MSA (24-31).

## Neuropathological characteristics and filament characterisation

We used sarkosyl to extract filaments from the cingulate cortex of an individual with a neuropathologically confirmed diagnosis of PD, two individuals with PDD and one individual with DLB (case 3). Frontal cortex was used for DLB cases 1 and 2. The individual with PD had a disease duration of 22 years and an age at death of 78 years; the individuals with PDD had disease durations of 8 and 13 years and ages at death of 87 and 76 years, respectively; DLB case 1 was in a 59-year-old individual with a disease duration of 10 years; DLB case 2 was in a 74-year-old individual with a disease duration of 13 years; DLB case 3 was in a 78-year-old individual with a disease duration of 15 years. Abundant Lewy bodies and Lewy neurites were stained by antibody Syn1, which is specific for the NAC region of α-synuclein (**Extended Data Figure 1**). Some glial cell staining was also present in DLB case 3. By negative-stain electron microscopy, cases of PD, PDD and DLB showed filaments with a diameter of 10 nm. Immunogold negative-stain electron microscopy with anti-α-synuclein antibody PER4 showed decoration of filaments, consistent with previous findings (**Extended Data Figure 2a-c**) (6,32,33). Immunoblotting of sarkosyl-insoluble material from the cases of PD, PDD and DLB with antibodies Syn303, Syn1 and PER4 showed high-molecular weight material (**Extended Data Figure 2d-f**). Full-length α-synuclein was the predominant species in most cases, but truncated α-synuclein was also present. The sequences of the coding exons of *SNCA* were wild-type in PD, PDD1, PDD2, DLB1, DLB2 and DLB3. We used cryo-EM to determine the atomic structures of α-synuclein filaments from all six cases (**Figure 1; Methods; Extended Data Figures 3 and 4; Extended Data Tables 1 and 2**).

**Figure 1.**
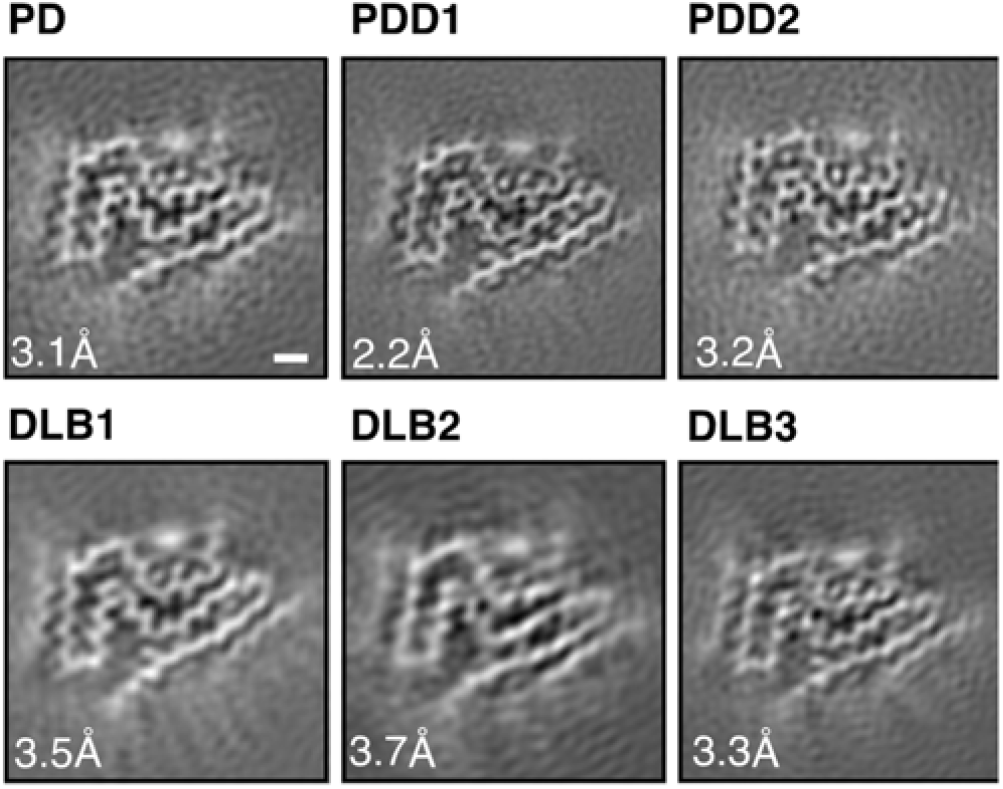
Cross-sections of α-synuclein filaments (Lewy fold) perpendicular to the helical axis, with a projected thickness of approximately one rung. PD, Parkinson’s disease; PDD, Parkinson’s disease dementia; DLB, Dementia with Lewy bodies. Scale bar, 1 nm.

## α-Synuclein filaments of PD, PDD and DLB

In agreement with previous observations for DLB (6), most filaments from the cases with PD, PDD and DLB did not exhibit a helical twist in the cryo-EM micrographs. Still, for each case, a minority of filaments (∼25%) was twisted, allowing their structure determination by helical reconstruction. α-Synuclein filaments from PD, PDD and DLB are identical and comprise a single protofilament (**Figures 1 and 2**). We termed the structure of the ordered core of these filaments “Lewy fold”. The reconstruction with the highest resolution, 2.2 Å for case 1 of PDD, showed density for main-chain oxygen atoms (**Extended Data Figure 3c**), establishing that α-synuclein filaments with the Lewy fold have a right-handed twist, in contrast to the left-handed twist observed for α-synuclein filaments with the MSA fold (6).

**Figure 2.**
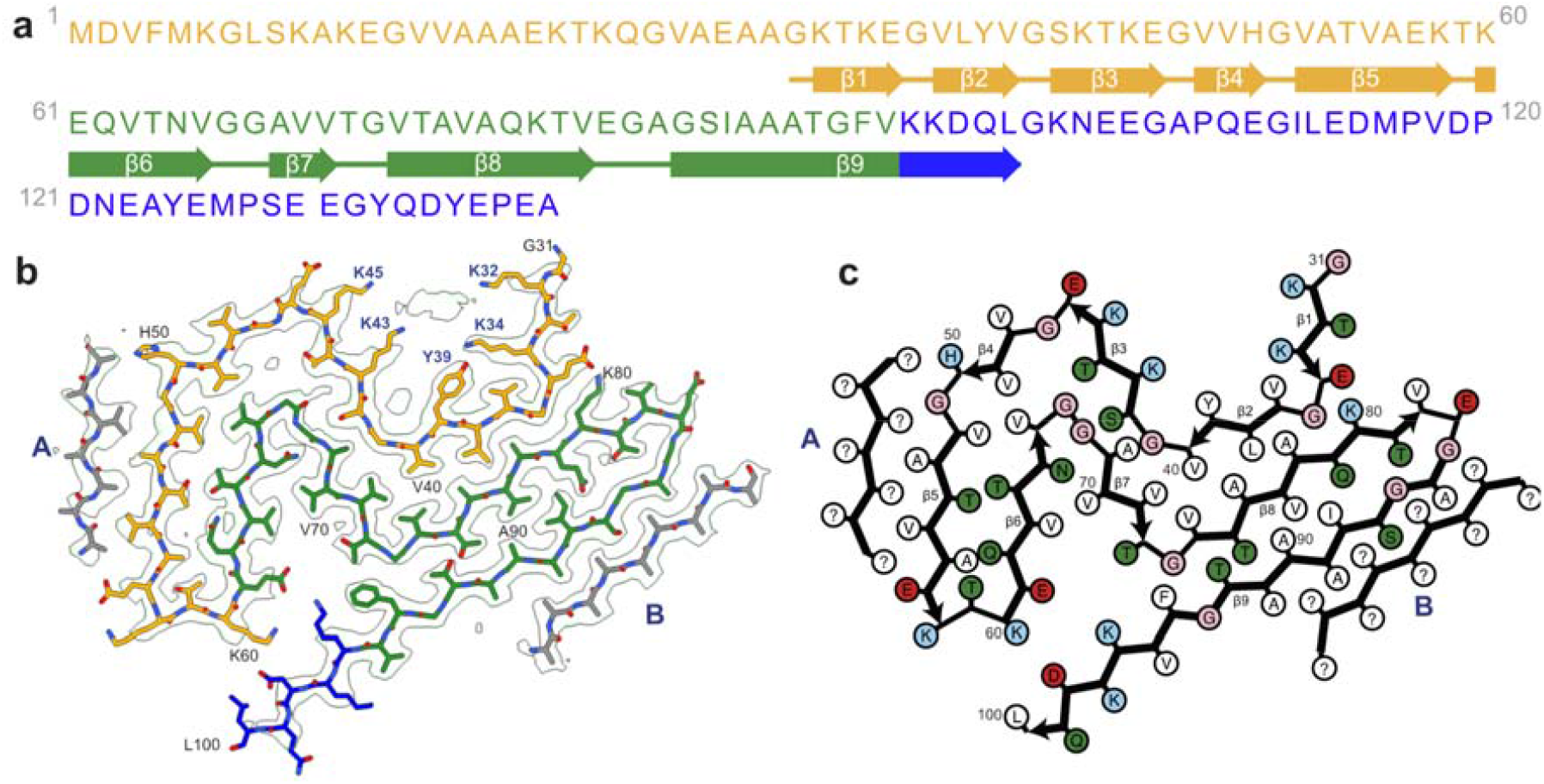
Cryo-EM structure of α-synuclein filaments from Parkinson’s disease, Parkinson’s disease dementia and dementia with Lewy bodies (Lewy fold). **(a)**. Amino acid sequence of human α-synuclein. N-terminal region (residues 1-60) in orange, NAC region (residues 61-95) in green and C-terminal region (residues 96-140) in blue. Thick connecting lines with arrowheads indicate β-strands. (**b)**. Cryo-EM density map and atomic model of the Lewy fold. The filament core extends from G31-L100. Islands A and B are indicated in grey. **(c)**. Schematic of the Lewy filament fold of α-synuclein. Negatively charged residues are in red, positively charged residues in blue, polar residues in green, apolar residues in white, sulfur-containing residues in yellow and glycines in pink. Thick connecting lines with arrowheads indicate β-strands. Unknown residues are indicated by question marks.

We did not determine the structures of the untwisted filaments. Nevertheless, most 2D class averages of untwisted filaments resembled projections of untwisted filament models with the Lewy fold (**Extended Data Figure 4a-h**). Moreover, stretches of segments that gave rise to twisted 2D class averages were typically observed together with stretches of segments giving rise to untwisted 2D class averages within the same filaments (**Extended Data Figure 4i-k**). It is thus likely that most of the untwisted filaments also adopted the Lewy fold, although we cannot exclude the presence of additional, minority folds among untwisted filaments. It is possible that cryo-EM grid preparation leads to untwisting of filaments, and it remains to be investigated whether filaments with a right-handed twist are more prone to untwisting than those with a left-handed twist.

The Lewy fold is formed by residues 31-100 of α-synuclein, which arrange as nine β-strands (β1-9) in a three-layered structure (**Figure 2**). The first two layers are corrugated, with the first comprising β1-5 and the second β6-8. The third layer consists only of β9. Two additional, partial layers are made by densities (islands) that are not connected to the rest of the ordered core. Island A packs against β5; island B packs against the N-terminal half of β9. The reconstructed densities for both islands indicate that they are made of peptides, but a lack of distinct side chain densities precluded their identification.

Besides the density for the ordered core of α-synuclein filaments and islands A and B, there are several additional densities in the cryo-EM reconstruction that could not be explained by peptides. We observed a density that is approximately 55 Å^3^ in size, in front of K32, K34, Y39, K43 and K45 (**Figure 3**). Its size and chemical environment resemble those of the unidentified cofactors in the α-synuclein filaments from MSA. Its position, in the middle of an outside groove formed by β1-3, suggests that the corresponding cofactors may be important for formation of the Lewy fold. Smaller, spherical densities next to Y39 and T44 probably corresponded to solvent molecules.

**Figure 3.**
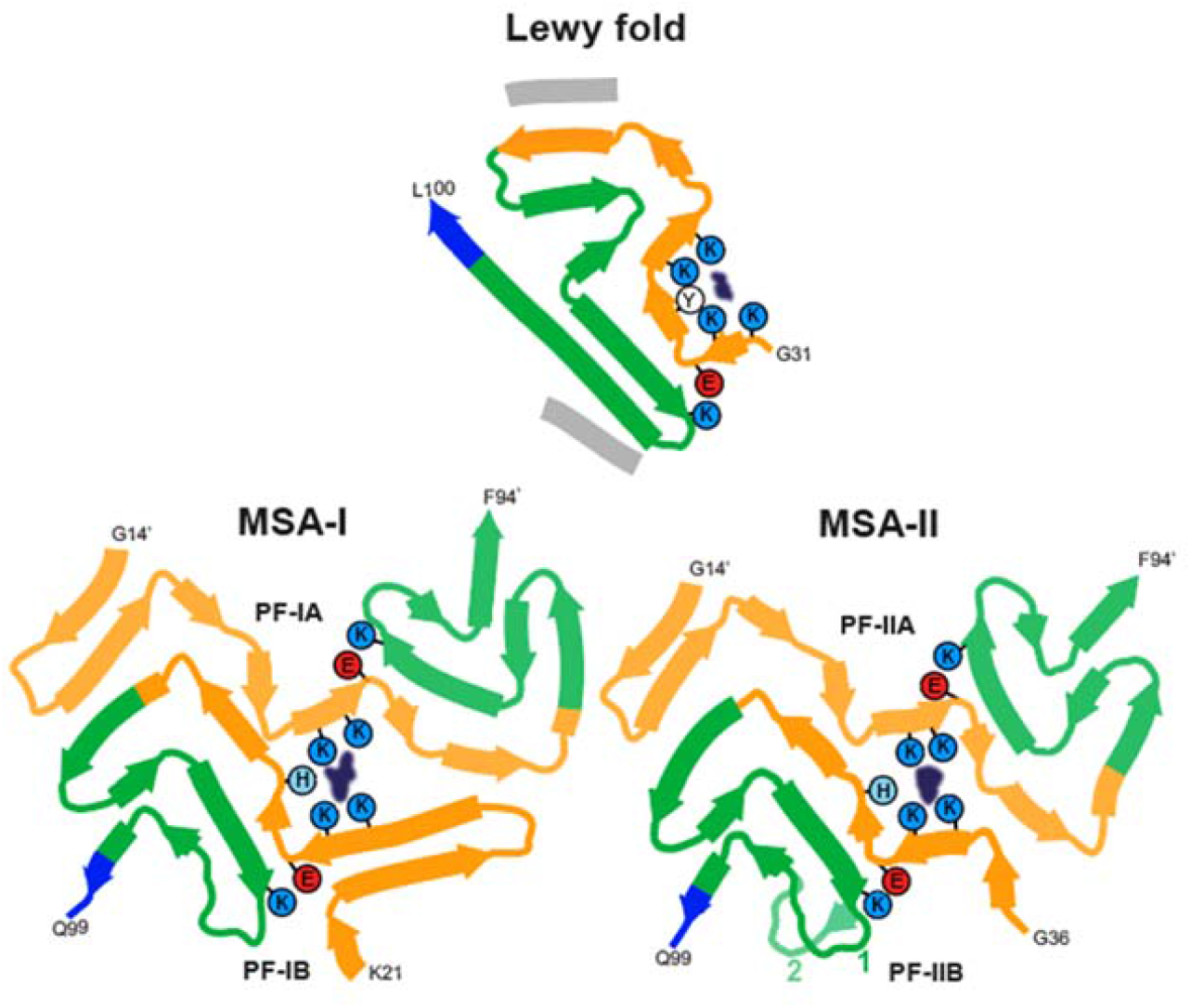
Comparison of the Lewy and MSA α-synuclein filament folds. Schematic of secondary structure elements in the Lewy and MSA folds, depicted as a single rung, and coloured as in Figure 2 (N-terminal region of α-synuclein in orange, NAC region in green and C-terminal region in blue; thick connecting lines with arrowheads indicate β-strands). The extra densities in all structures are depicted in dark blue. The positions of their surrounding residues, as well as the supporting salt bridges between E35 and K80 in the Lewy fold and between E46 and K80 in MSA protofilaments, are highlighted with coloured circles.

The Lewy fold differs from the structures of MSA filaments from human brains (6) and from those of *in vitro* assembled α-synuclein filaments (34). Whereas Lewy and MSA folds are different, substructures of the Lewy fold have been observed in α-synuclein filaments that were assembled *in vitro* (**Figure 3; Extended Data Figure 5**). Residues 32-41 and 70-82 overlay with a root-mean-square deviation of the main-chain atoms (rmsd) of 1.1 Å with the same residues in a recombinant α-synuclein filament with phosphorylation of Y39 (35) (PDB:6L1T). Residues 42-67 also overlay with a rmsd of 1.3 Å onto the same structure, albeit in a different orientation relative to the rest of the fold. Both substructures contribute to the electrostatic interaction network around the phosphate group of Y39. However, in the Lewy fold, there is no additional density at Y39, indicating that this residue is not phosphorylated. Instead, both substructures form the cofactor-binding groove. In the “hinge” region, between residues 42-67 and 70-82, residues 61-72 adopt essentially the same conformation as in the α-synuclein filaments from MSA and some filaments assembled from recombinant proteins. In addition, residues 69-98 of the Lewy fold overlay with a rmsd of 0.5 Å with the same residues in filaments of N-terminally truncated recombinant α-synuclein (41-140) (36) (PDB:7LC9). Interestingly, residues 85-92, as well as the peptide corresponding to the density of island A, overlay with a rmsd of 0.5 Å with residues 14-23 and 85-92 of recombinant α-synuclein filament polymorph 2a that was assembled from full-length recombinant protein (36) (PDB:6SSX). In this structure, the density for residues 14-23 is also disconnected from the rest of the ordered core. It is therefore likely that the peptide corresponding to island A in the Lewy fold corresponds to part of the α-synuclein sequence that is N-terminal of the ordered core. The close packing of island A against β9 of the Lewy fold suggests that the corresponding interface comprises at least two consecutive small residues, which pack in the spaces between A85, S87 and A89 of β9. Such residues are present in the α-synuclein sequence that is N-terminal to the ordered core, e.g., G7, S9, A11, A17, A19, A27, A29, but not in the C-terminus. Although residues 14-23 of α-synuclein fit into the island A density, for them to be part of the same protein chain, the (presumably disordered) connecting residues must adopt a fully extended conformation. In polymorph 2a, there is also a second substructure (residues 52-66) with similarity to the Lewy fold; it is a shorter segment than in the corresponding substructure of the recombinant α-synuclein filament with phosphorylation of Y39.

Residues 47-60 of α-synuclein, together with the peptide responsible for the density of island B, resemble the dimeric interface found in several α-synuclein filaments formed from recombinant proteins *in vitr*o, for instance polymorph 1a (PDB:6H6B) (37), overlaying residues 47-60 from one protofilament with residues 50-57 from the other, with a rmsd of 0.9 Å. The island B interface of the ordered core harbours mutations that cause inherited PD, such as G51D and A53E/G/T/V (2). Most mutations are incompatible with the interfaces of filaments assembled *in vitro*, suggesting that they are also likely to disrupt the interface with island B peptides.

## Implications

We establish the existence of molecular conformers of assembled α-synuclein in neurodegenerative disease (**Figure 3**). For tau, distinct conformers define different diseases (38). Lewy body diseases PD, PDD and DLB share the same protofilament fold, confirming that they are closely related. Dementia is common in PD, especially in advanced cases (1-3). A diagnosis of PDD is made when cognitive impairment develops in a patient with long-standing idiopathic PD, whereas dementia that develops within a year of PD is called DLB (1,4). PDD and DLB show similar neuropathological profiles, including the presence of widespread cortical α-synuclein Lewy pathology (39). Many cases have also some Alzheimer-type plaques and tangles. The combination of Lewy body- and Alzheimer-type pathologies correlates well with PDD and DLB (1,40). Consistent with the presence of the same protofilament in PD, PDD and DLB, Lewy pathology in the brain forms first in the brainstem, from where it progresses to limbic and cortical areas (41). These findings indicate that PD, PDD and DLB are part of a continuum of diseases. Lewy pathology is also characteristic of incidental Lewy body disease, primary autonomic failure, many cases of rapid eye movement sleep behaviour disorder and some cases of Alzheimer’s disease (1,2). It remains to be seen if the α-synuclein filament structures are the same as those reported here. This is also true of the Lewy pathology found in the peripheral nervous system in PD.

The filament fold of Lewy body diseases differs from that of MSA, consistent with seed amplification being able to distinguish between PD and MSA (30,42). However, known structures of seeded α-synuclein aggregates are different from those of the seeds (43,44). Unlike for Lewy body diseases, two filament types have been observed in MSA, each of which is made of two different protofilaments (6). The differences between Lewy and MSA folds are consistent with differences in morphology that were described between the α-synuclein filaments of DLB and MSA by negative staining (45). In the α-synuclein filament structures of MSA, E46 forms a salt bridge with K80, whereas E35 forms a salt bridge with K80 in PD, PDD and DLB (**Figure 3)** This may explain why DLB seeds, unlike those from MSA, induced the seeded assembly of E46K α-synuclein in HEK cells (31,46).

Why are the Lewy and MSA folds of α-synuclein different? It will be important to know more about post-translational modifications and the identities of non-proteinaceous densities associated with these folds. Different cellular environments may play a role (23). Filaments of recombinant α-synuclein, including those amplified from brain seeds, failed to adopt the same fold as filaments from human brains. To understand the mechanisms that lead to the formation of α-synuclein folds, it is important to develop methods by which to assemble recombinant α-synuclein into Lewy and MSA folds, similar to what has been done for Alzheimer tau filaments (47). It will also be important to identify conditions for making seeded aggregates with structures identical to those of α-synuclein seeds and to produce animal models with structures of α-synuclein filaments like those from human brain. Knowledge of the structures of α-synuclein filaments and how they form may be used for the development of specific biomarkers for synucleinopathies and the development of safe and effective mechanism-based therapies.

## Acknowledgements

We thank the patients’ families for donating brain tissues, T. Darling and J. Grimmett for help with high-performance computing and the EM facility of the Medical Research Council (MRC) Laboratory of Molecular Biology for help with cryo-EM data acquisition. We thank R.A. Crowther, S. Lövestam, W. Poewe, M.G. Spillantini and E. Tolosa for helpful discussions. We acknowledge Diamond Light Source for access and support of the cryo-EM facilities at the UK’s Electron Bio-imaging Centre (under proposal bi23268), funded by the Wellcome Trust, the MRC and the Biotechnology and Biological Sciences Research Council (BBSRC). This work was supported by the MRC (MC_UP_A025_1013 to S.H.W.S. and MC_U105184291 to M.G.). T.L. holds an Alzheimer’s Research UK Senior Fellowship. T.R. is supported by the National Institute for Health Research Queen Square Biomedical Research Unit in Dementia. The Queen Square Brain Bank is supported by the Rita Lila Weston Institute for Neurological Studies. This work was also supported by the Japan Agency for Science and Technology (CREST) (JPMJCR18H3 to M.H.), the Japan Agency for Medical Research and Development (AMED) (JP20dm0207072 to M.H.), the US National Institutes of Health (P30-AG010133, U01-NS110437 and RF1-AG071177, to R.V. and B.G., and R01NS037167, to T.F.) and the Department of Pathology and Laboratory Medicine, Indiana University School of Medicine (to R.V. and B.G.).

## Author Contributions

P.W.C., Yuko Saito, T.F., T.T.W., K.H., S.M., T.R., B.G., M.H. and T.L. identified patients and performed neuropathology; Y.Y., H.J.G., R.V. and M.H. performed analysis of brain samples; Y.Y., Yang Shi, M.S., X.Z. and A.K. collected cryo-EM data; Y.Y., Yang Shi, M.S., A.G.M. and S.H.W.S. analysed cryo-EM data; Y.Y. performed immunoblot analysis; B.G., M.H. and T.L. performed immunohistochemistry; S.H.W.S. and M.G. supervised the project. All authors contributed to the writing of the manuscript.

## Additional Information

For the purpose of open access, the MRC Laboratory of Molecular Biology has applied a CC BY public copyright licence to any Author Accepted Manuscript version arising.

## Data Availability

Cryo-EM maps have been deposited in the Electron Microscopy Data Bank (EMDB) under accession number 15285. Corresponding refined atomic models have been deposited in the Protein Data Bank (PDB) under accession number 8A9L.

## Methods

No statistical methods were used to predetermine sample size. The experiments were not randomized and investigators were not blinded to allocation during experiments and outcome assessment.

### Clinical history and neuropathology

PD was in a 78-year-old woman who died with a neuropathologically confirmed diagnosis after a 22-year history of slowly progressing rest tremor and bradykinesia. PDD case 1 was in an 87-year-old man who died with a neuropathologically confirmed diagnosis following an 8-year history of prominent rest tremor and bradykinesia. He developed dementia approximately 3 years after the diagnosis of PD. This case has been described before [case 12 in (48)]. PDD case 2 was in a 76-year-old woman who died with a neuropathologically confirmed diagnosis after a 13-year history of disturbed sleep, orthostatic hypotension, resting tremor and bradykinesia. She began to develop dementia approximately 4 years after the diagnosis of PD. The individuals with DLB have been described before (6). They developed dementia around the same time as PD.

### Sequencing of *SNCA* coding exons

Genomic DNA was extracted from frozen brain tissues. Coding exons of *SNCA* and flanking intronic sequences were amplified by polymerase chain reaction and sequenced using the dideoxy method.

### Extraction of α-synuclein filaments

Sarkosyl-insoluble material was extracted from fresh-frozen cingulate cortex and frontal cortex of individuals with PD, PDD and DLB, essentially as described (25). In brief, tissues were homogenized in 20 vol (v/w) extraction buffer consisting of 10 mM Tris-HCl, pH 7.5, 0.8 M NaCl, 10% sucrose and 1 mM EGTA. Homogenates were brought to 2% sarkosyl and incubated for 30 min at 37° C. Following a 10 min centrifugation at 10,000g, the supernatants were spun at 100,000g for 20 min. Pellets were resuspended in 500 µl/g extraction buffer and centrifuged at 3,000g for 5 min. Supernatants were diluted threefold in 50 mM Tris-HCl, pH 7.5, containing 0.15 M NaCl, 10% sucrose and 0.2% sarkosyl, and spun at 166,000g for 30 min. Sarkosyl-insoluble pellets were resuspended in 100 µl/g of 20 mM Tris-HCl, pH 7.4.

### Immunolabelling and histology

Immunogold negative-stain electron microscopy and immunoblotting were carried out as described (49). Filaments were extracted from cingulate cortex of the case of PD, PDD cases 1 and 2, as well as DLB case 3. Frontal cortex was used for DLB cases 1 and 2. PER4, a rabbit polyclonal serum that was raised against a peptide corresponding to residues 116-131 of human α-synuclein (32), was used at 1:50. Images were acquired at 11,000x with a Gatan Orius SC200B CCD detector on a Tecnai G2 Spirit at 120 kV. For immunoblotting, samples were resolved on 4-12% Bis-Tris gels (NuPage) and primary antibodies diluted in PBS plus 0.1% Tween 20 and 5% non-fat dry milk. Before blocking, membranes were fixed with 1% paraformaldehyde for 30 min. Primary antibodies were: Syn303 [a mouse monoclonal antibody that recognizes residues 1-5 of human α-synuclein (50)] (BioLegend) at 1:4,000, Syn1 [a mouse monoclonal antibody that recognizes residues 91-99 from the NAC region of human α-synuclein (51)] (BD Biosciences) at 1:4,000 and PER4 at 1:4,000. Histology and immunohistochemistry were carried out as described (25,52). Some sections (8 µm) were counterstained with haematoxylin. The primary antibody was Syn1 (1:1,000).

### Electron cryo-microscopy

Extracted filaments were centrifuged at 3,000 g for 3 min and applied to glow-discharged holey carbon gold grids (Quantifoil Au R1.2/1.3, 300 mesh), which were glow-discharged with an Edwards (S150B) sputter coater at 30 mA for 30 s. Aliquots of 3 μl were applied to the grids and blotted with filter paper (Whatman, cat no. 1001-070) at 100% humidity and 4°C, using a Vitrobot Mark IV (Thermo Fisher Scientific). For all cases, datasets were acquired on Titan Krios G2, G3 and G4 microscopes (Thermo Fisher Scientific) operated at 300 kV. Images for the case of PD and case 2 of PDD were acquired using a Falcon-4 detector (Thermo Fisher Scientific). Images for case 1 of PDD and case 1 of DLB were acquired using a Falcon-4i detector (Thermo Fisher Scientific) in EER mode with a flux of 8 electrons/pixel/s and a Selectris-X energy filter (Thermo Fisher Scientific) with a slit width of 10 eV to remove inelastically scattered electrons. Images for cases 1-3 of DLB were acquired with a Gatan K2 or K3 detector in super-resolution counting mode, using a Bio-quantum energy filter (Gatan) with a slit width of 20 eV. Images were recorded with a total dose of 40 electrons per Å^2^.

### Helical reconstruction

Movie frames were gain-corrected, aligned, dose-weighted and then summed into a single micrograph using RELION’s own motion correction program (53). Contrast transfer function (CTF) parameters were estimated using CTFFIND-4.1 (54). All subsequent image-processing steps were performed using helical reconstruction methods in RELION (55,56). α-Synuclein filaments were picked manually, as they could be distinguished from filaments made of tau, Aβ, and TMEM106B by their general appearance (**Extended Data Table 1**). For all data sets, reference-free 2D classification was performed to select suitable segments for further processing. For the case of PD, PDD case 2 and DLB case 2, start-end coordinates for twisting α-synuclein filaments were re-picked manually, based on individual segments that were assigned to 2D classes that corresponded to twisting filaments. Initial 3D reference models were generated *de novo* from 2D class averages using an estimated rise of 4.75 Å and helical twists according to the observed cross-over distances of the filaments in the micrographs (57) for PDD case 1. The refined model from PDD case 1, low-pass filtered to 10 Å, was used as initial model for the other cases of PD, PDD and DLB. Combinations of 3D auto-refinements and 3D classifications were used to select the best segments for each structure. For all data sets, Bayesian polishing (52) and CTF refinement (58) were performed to further increase the resolution of the reconstructions. Final reconstructions were sharpened using standard post-processing procedures in RELION, and overall final resolutions were estimated from Fourier shell correlations at 0.143 between the independently refined half-maps, using phase randomisation to correct for convolution effects of a generous, soft-edged solvent mask (59) (**Extended Data Figure 3**). Untwisted models with the Lewy fold and their projections were generated using the relion helix inimodel2d and relion project programs, respectively.

### Model building

Atomic models comprising three β-sheet rungs were built *de novo* in Coot (60) in the best available map for PDD case 1. Coordinate refinements were performed using *Servalcat* (61). Final models were obtained using refinement of only the asymmetric unit against the half-maps in *Servalcat*.

## EXTENDED DATA FIGURES

**Extended Data Figure 1.**
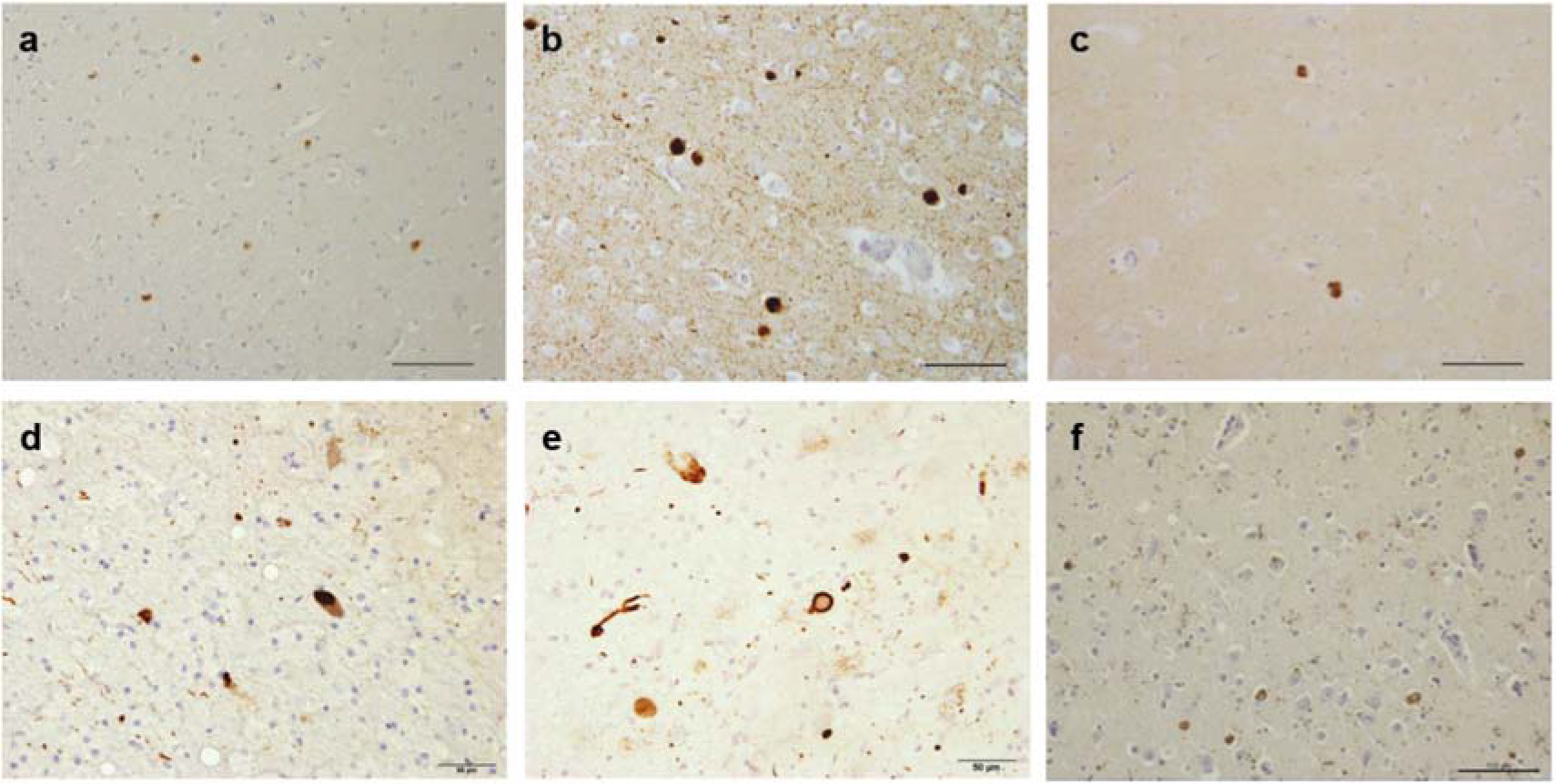
Immunostaining of α-synuclein inclusions. Sections from brain regions contralateral to those used for cryo-EM structure determination were stained with monoclonal antibody Syn1 (1:1,000). (a), Cingulate cortex from PD; (b), Cingulate cortex from PDD1; (c), Cingulate cortex from PDD2; (d), Frontal cortex from DLB1; (e), Frontal cortex from DLB2; (f), Cingulate cortex from DLB3. Scale bars: a-c, f, 100 μm; d,e, 50 μm.

**Extended Data Figure 2.**
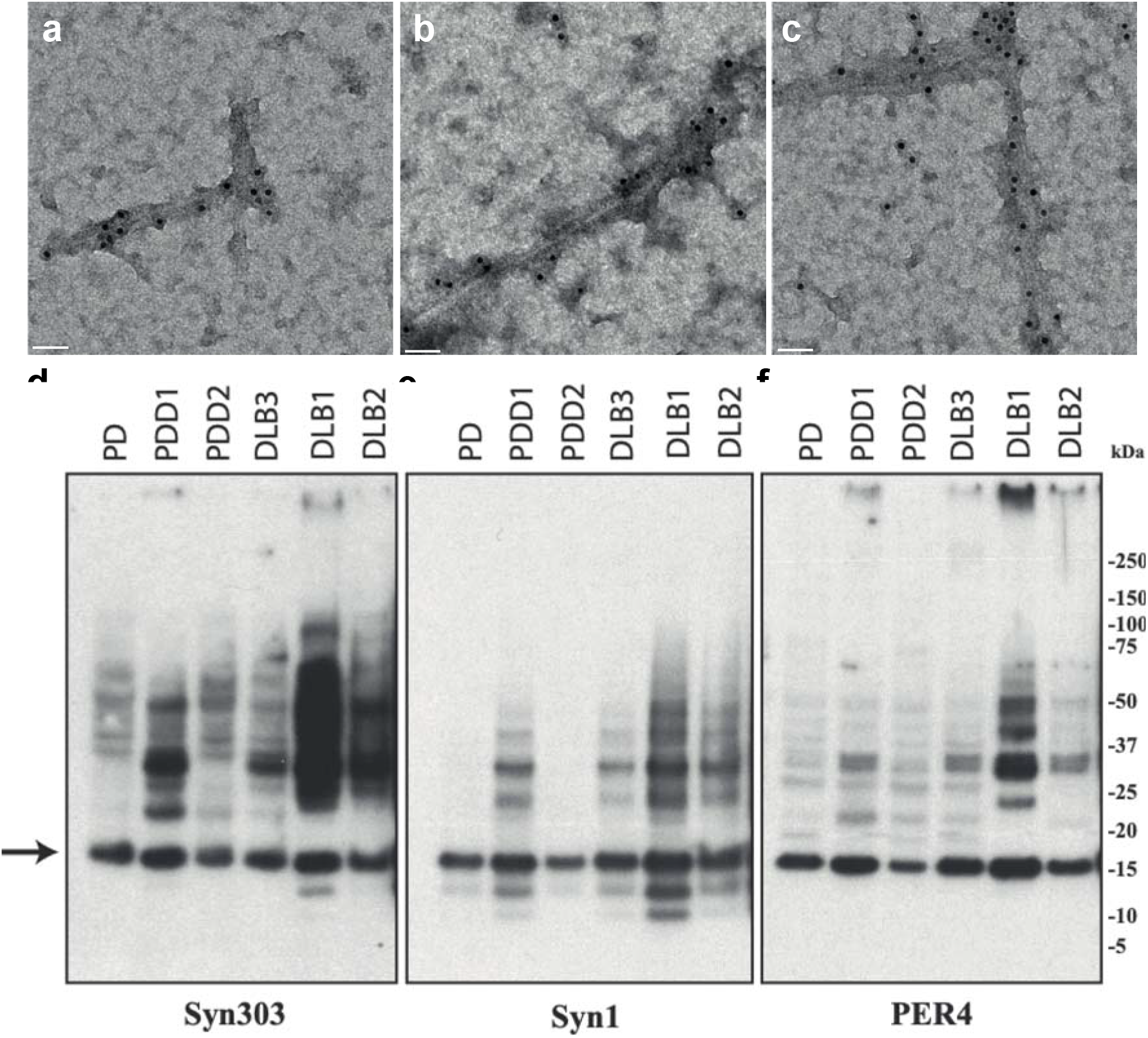
Negative-stain immunoelectron microscopy and immunoblotting of sarkosyl-insoluble material. PER4 was used at 1:50 in (a-c). (a), PD (Cingulate cortex); (b), PDD1 (Cingulate cortex); (c), DLB3 (Cingulate cortex); Syn303, Syn1 and PER4 were used at 1:4,000 in (d-f). The brain regions used for cryo-EM were also used for immunoblotting. The arrow points to the position of monomeric α-synuclein.

**Extended Data Figure 3.**
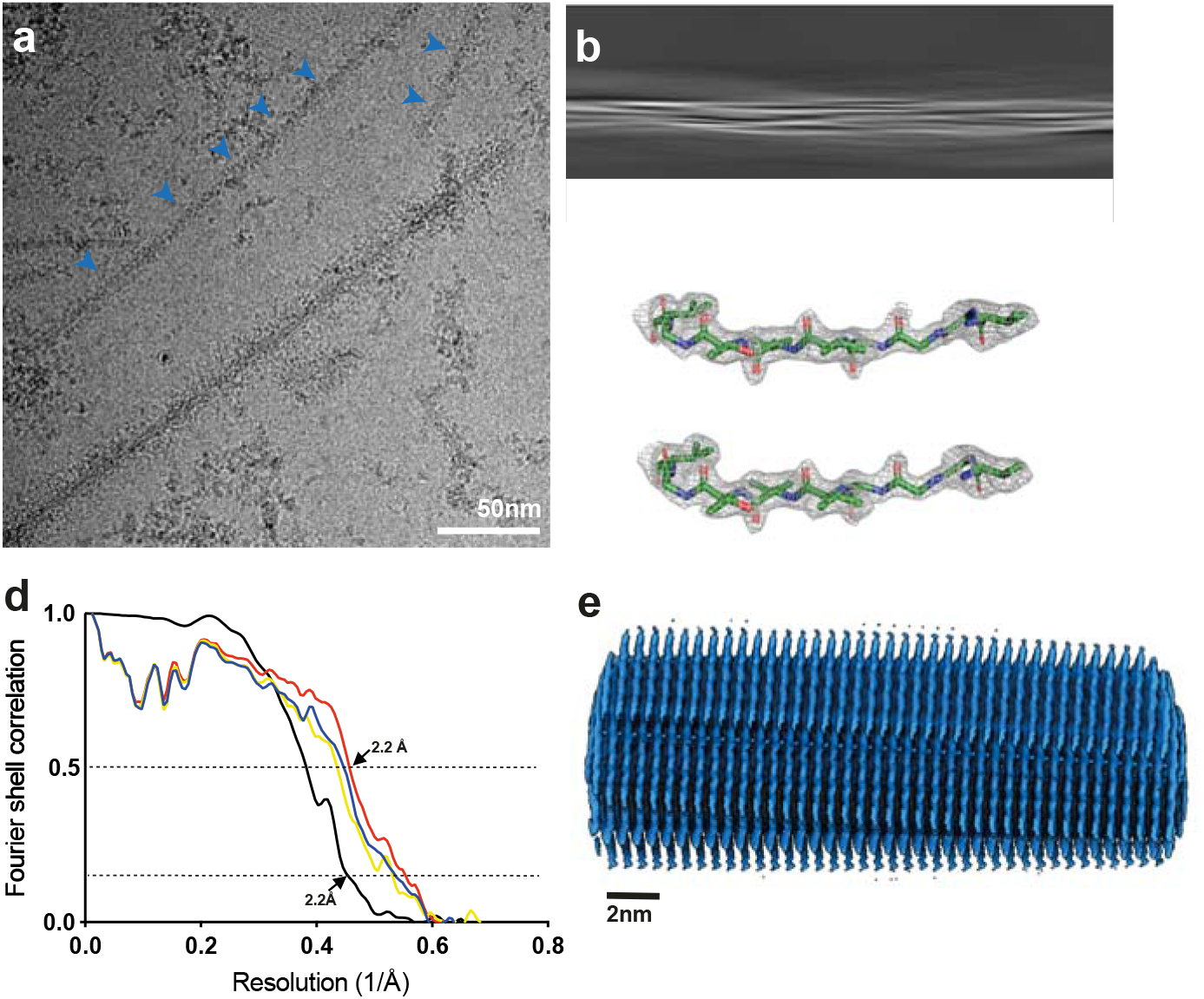
Cryo-EM maps, cryo-EM images and resolution estimates. (a), α-Synuclein filaments (blue arrows) from PDD1. Scale bars, 50 nm. (b), Projection features of Lewy filament. Scale bars, 5 nm. (c), Zoomed-in view of the main chain showing density of the oxygen atoms. (d), Fourier shell correlation (FSC) curves for cryo-EM maps are shown in black; for the final refined atomic model against the final cryo-EM map in red; for the atomic model refined in the first half map against that half map in blue; for the refined atomic model in the first half map against the other half map in yellow. (e), Side view of the Lewy fold.

**Extended Data Figure 4.**
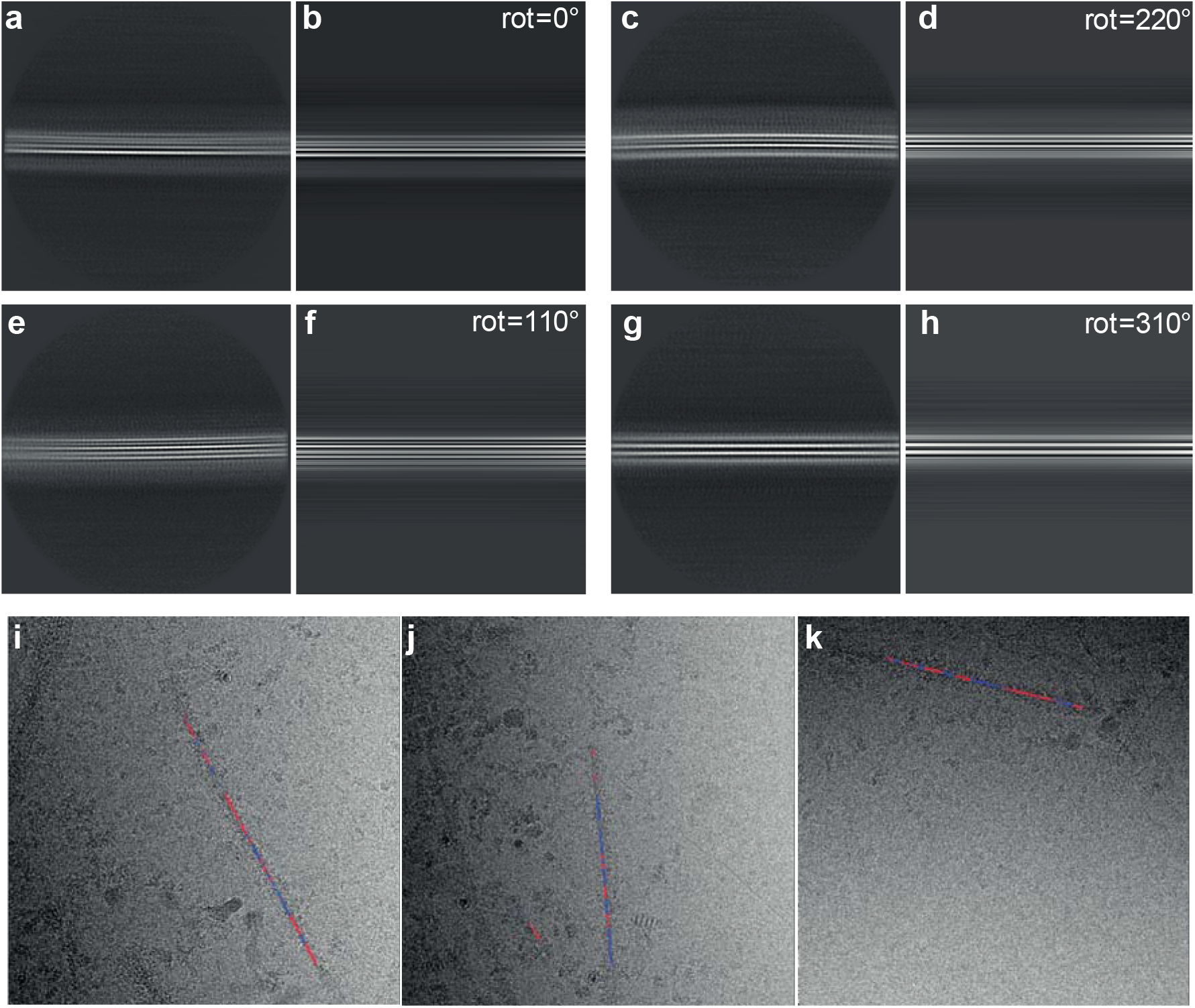
Twisted and untwisted filaments in 2D class averages and micrographs. Case 1 of PDD was used. (a,c,e,g), 2D class averages of untwisted filaments; (b,d,f,h) projections of untwisted models with the Lewy fold, rotated by 0, 220, 110 and 310 degrees, respectively, along the first Euler angle (rot). Box size, 640 Å. (i,j,k), Micrographs of untwisted and twisted filaments. Blue indicates segments that contributed to twisted 2D class averages and red segments that contributed to untwisted 2D class averages.

**Extended Data Figure 5.**
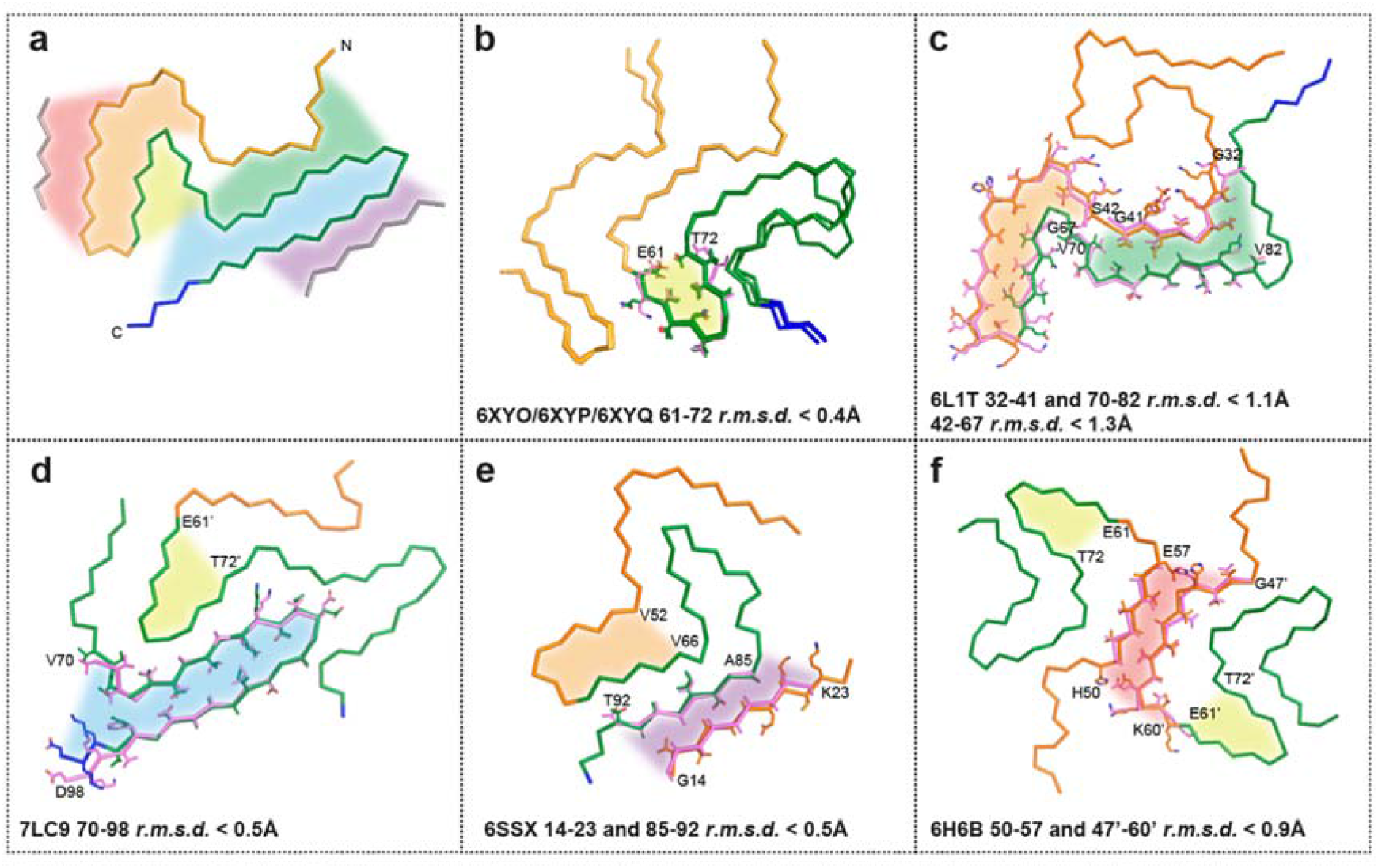
Comparison of the Lewy fold with structures of α-synuclein filaments from human brains or assembled from recombinant proteins. (a), Ribbon plot of the Lewy fold; the protein chain is coloured as in Figure 2. Highlighted by red, orange, yellow, green, blue and purple areas are substructures that are individually shared with other filament structures. These local similarities are indicated with the same coloured areas and the overlays of the corresponding substructures are shown in sticks on the following panels (b-f). (b), Common core structure of MSA Type I and Type II filaments (made of PFIA/IIA 14-47 and PFIB/IIB 41-99), with a shared substructure highlighted in yellow. (c), pY39 α-synuclein protofilament (PDB:6L1T) with two different substructures highlighted in orange and green. (d), N-terminally truncated α-synuclein (40-140) dimeric filament (PDB:7LC9), with two different substructures in its protofilaments, highlighted in blue and yellow. (e), Polymorph 2a filament (PDB:6SSX), with two substructures highlighted in purple and orange. (f), Polymorph 1a filament (PDB:6H6B) contains yellow-coloured substructures in its protofilaments and a red-coloured substructure in their dimeric interface.

## EXTENDED DATA TABLES

**Extended Data Table 1.** Filament types.

| Case | Disease | Age (yrs) | α-Synuclein | Tau | Aβ42 | TMEM106B |
| --- | --- | --- | --- | --- | --- | --- |
| 1 | PD | 78 | 55% | <1% | 10% | 35% |
| 2 | PDD1 | 87 | 42% | 38% | 5% | 15% |
| 3 | PDD2 | 76 | 58% | <1% | 5% | 37% |
| 4 | DLB1 | 59 | 60% | 10% | 27% | 3% |
| 5 | DLB2 | 74 | 59% | <1% | 5% | 36% |
| 6 | DLB3 | 78 | 54% | 12% | 22% | 12% |

**Extended Data Table 2.**
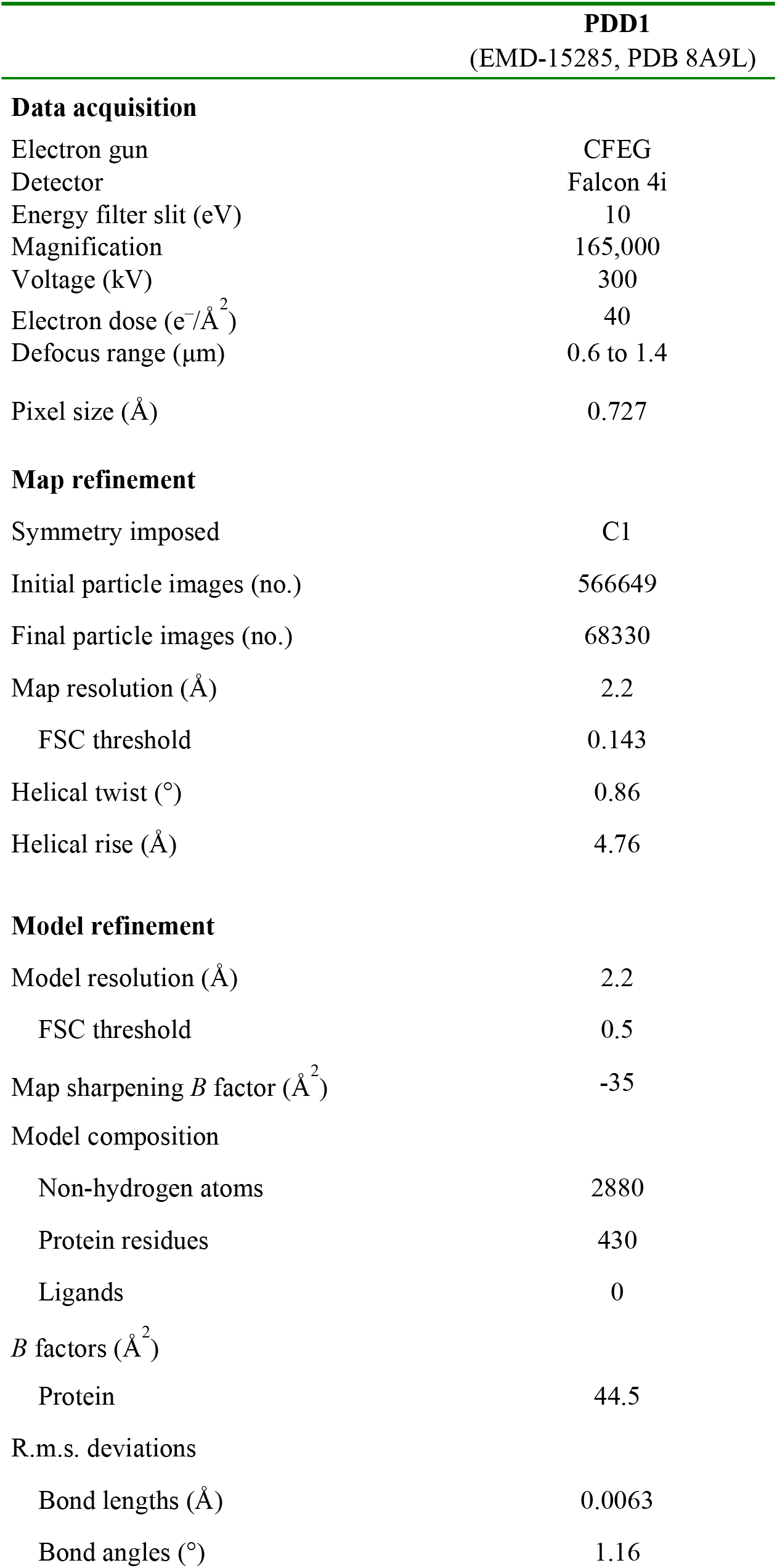
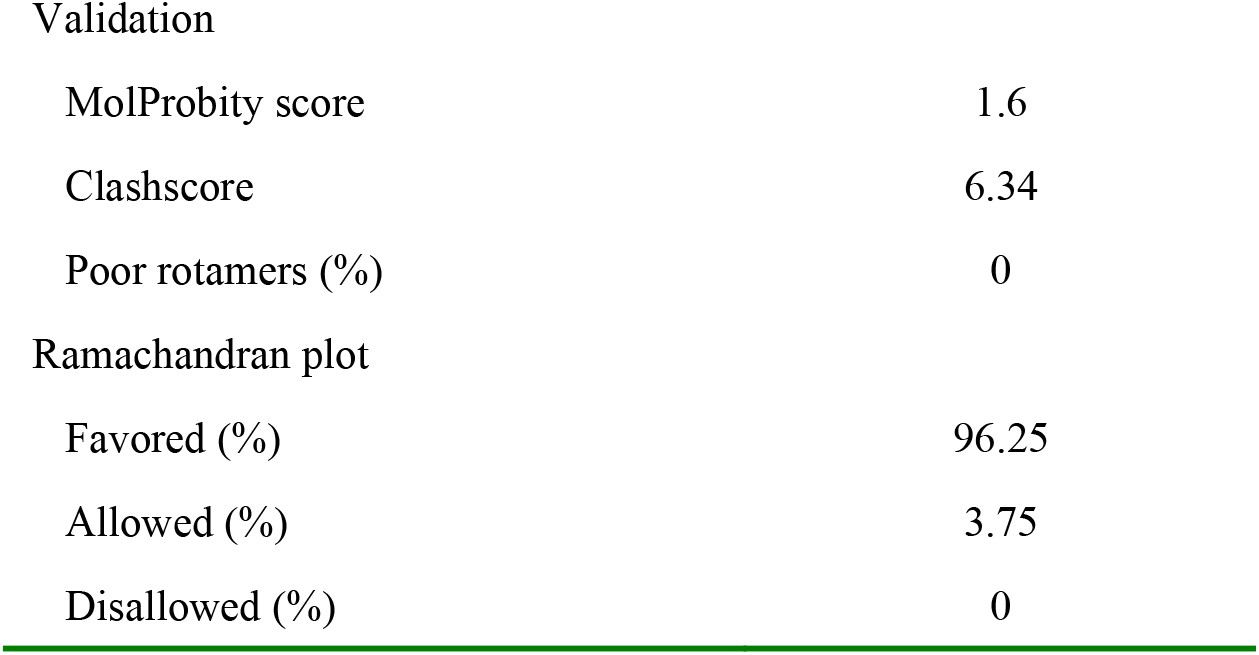
Cryo-EM data acquisition and structure determination.

| **PDD1**<br>(EMD-15285, PDB 8A9L) |  |
| --- | --- |
| **Data acquisition** |  |
| Electron gun | CFEG |
| Detector | Falcon 4i |
| Energy filter slit (eV) | 10 |
| Magnification | 165,000 |
| Voltage (kV) | 300 |
| Electron dose ($\text{e}^-/\text{\AA}^2$) | 40 |
| Defocus range ($\mu\text{m}$) | 0.6 to 1.4 |
| Pixel size ($\text{\AA}$) | 0.727 |
| **Map refinement** |  |
| Symmetry imposed | C1 |
| Initial particle images (no.) | 566649 |
| Final particle images (no.) | 68330 |
| Map resolution ($\text{\AA}$) | 2.2 |
| FSC threshold | 0.143 |
| Helical twist ($^\circ$) | 0.86 |
| Helical rise ($\text{\AA}$) | 4.76 |
| **Model refinement** |  |
| Model resolution ($\text{\AA}$) | 2.2 |
| FSC threshold | 0.5 |
| Map sharpening $B$ factor ($\text{\AA}^2$) | -35 |
| Model composition |  |
| Non-hydrogen atoms | 2880 |
| Protein residues | 430 |
| Ligands | 0 |
| $B$ factors ($\text{\AA}^2$) | |
| Protein | 44.5 |
| R.m.s. deviations |  |
| Bond lengths ($\text{\AA}$) | 0.0063 |
| Bond angles ($^\circ$) | 1.16 |
| Validation |  |
| MolProbity score | 1.6 |
| Clashscore | 6.34 |
| Poor rotamers (%) | 0 |
| Ramachandran plot |  |
| Favored (%) | 96.25 |
| Allowed (%) | 3.75 |
| Disallowed (%) | 0 |

